# The FunCoup Cytoscape App: multi-species network analysis and visualization

**DOI:** 10.1101/2024.10.04.616627

**Authors:** Davide Buzzao, Lukas Steininger, Dimitri Guala, Erik L.L. Sonnhammer

**Affiliations:** Department of Biochemistry and Biophysics, Stockholm University, Science for Life Laboratory, Box 1031, 171 21 Solna, Sweden

## Abstract

**Summary:** Functional association networks, such as FunCoup, are crucial for analyzing complex gene interactions. To facilitate the analysis and visualization of such genome-wide networks, there is a need for seamless integration with powerful network analysis tools like Cytoscape.

The FunCoup Cytoscape App integrates the FunCoup web service API with Cytoscape, allowing users to visualize and analyze gene interaction networks for 640 species. Users can input gene identifiers and customize search parameters, employing various network expansion algorithms like group or independent gene search, MaxLink, and TOPAS. The app maintains consistent visualizations with the FunCoup website, providing detailed node and link information, including tissue and pathway gene annotations. The integration with Cytoscape plugins, such as ClusterMaker2, enhances the analytical capabilities of FunCoup, as exemplified by the identification of the *Myasthenia gravis* disease module along with potential new therapeutic targets.

**Availability and implementation:** The FunCoup Cytoscape App is developed using the Java OSGi framework, with UI components implemented in Java Swing and build support from Maven. The App is available as a JAR file at https://bitbucket.org/sonnhammergroup/funcoup_cytoscape/ repo, and can be downloaded from the Cytoscape App store https://apps.cytoscape.org/.

## INTRODUCTION

Understanding gene and protein interactions is crucial for deciphering complex cellular mechanisms. Modern high-throughput biotechnologies continuously generate vast amounts of new data, and to make sense of this data it is essential to interpret it within the cellular context and integrate existing biological knowledge. One of the most established resources for this purpose is FunCoup, which integrates diverse types of data to provide insights into functional associations between genes/proteins.

FunCoup is notable for its comprehensiveness, which is attributable to its unique bin-free redundancy-weighted naïve Bayesian training, combining ten diverse data types and extensive use of orthology transfer between species (See Figure 1). FunCoup release 6 (Buzzao et al. 2024) has increased coverage further, and features an improved web user interface for efficient network analysis and visualization. It extends the number of species to 640, uses a new training framework, and contains a large amount of directed regulatory interactions. FunCoup 6 further enhances the network search by incorporating the disease module algorithms TOPAS (Buzzao et al. 2022) and MaxLink (Guala, Sjölund, and Sonnhammer 2014). Additionally, FunCoup allows users to query multiple species simultaneously and perform comparative interactomics with customized network search algorithms.

**Figure 1.**
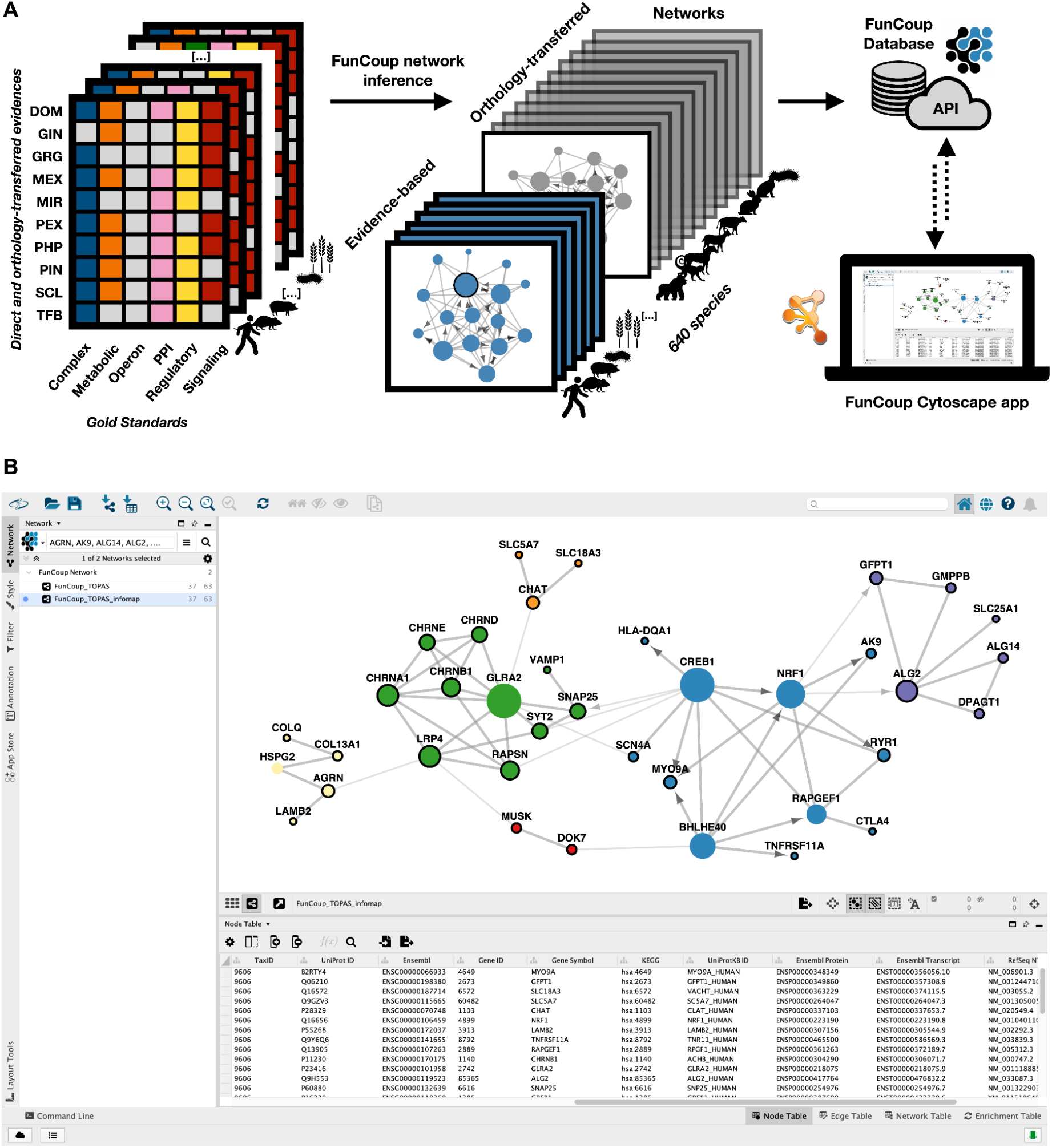
Overview of FunCoup and a use case. (A) FunCoup is a comprehensive resource of genome-wide functional association networks. FunCoup 6 uses six gold standards and ten different evidences to extract networks for 640 species. FunCoup is stored in a PostgreSQL database, and the FunCoup Cytoscape App connects the user interface to the RESTFUL API of the website https://funcoup6.scilifelab.se/. (B) Example from the FunCoup Cytoscape App. 31 genes known to be associated with *Myasthenia gravis* (Supplementary Table 2) were used as query/seed genes, and TOPAS was employed as search method, which added connector genes to produce a single disease module. The different node colors correspond to Infomap (ClusterMaker2) subclusters of the resulting disease module. Query/seed genes have a black border, while the found connector genes have none. The size of a node is proportional to its degree of connectivity. The link width is set by ClusterMaker2 to be thicker within clusters and thinner between clusters. Directions are shown for links with regulatory link score (GRG_LLR) > 1.

The Cytoscape (Shannon et al. 2003) community is very active with thousands of users and many developers. Over the years, Cytoscape has evolved into a rich environment with a large number of tools and plugins for network analysis and visualization. To reach this community, GeneMANIA and STRING previously developed Cytoscape plugins (Montojo et al. 2010; Doncheva et al. 2023). As an addition to the Cytoscape network database repertoire we here present the FunCoup Cytoscape App that retrieves data directly from the FunCoup web service APIs, bringing the FunCoup network resource into the Cytoscape ecosystem. In this way, the App enables users to utilize the extensive array of Cytoscape plugins and tools for advanced network analysis and visualization with the FunCoup networks. This direct connection allows researchers to make full use of the rich interaction data provided by FunCoup, improving their ability to conduct comprehensive and in-depth analyses within the powerful Cytoscape environment. To demonstrate this integration, we showcase an analysis of the network of *Myasthenia gravis* genes in combination with disease module detection and subclustering (Figure 1).

## RESULTS

The FunCoup App was developed to directly connect Cytoscape to the FunCoup web service API. The app replicates the visualizations found on the FunCoup website, providing a familiar and robust interface for users. In addition to FunCoup’s own features, the app allows users to access the extensive network analysis and visualization tools available in Cytoscape, making it possible to fully leverage the rich data available from FunCoup.

The FunCoup API can be queried for 640 species, 22 of which have evidence-based networks and the remaining 618 have orthology-transferred networks. The query process in the Cytoscape app is designed to handle complex data inputs efficiently. Users begin by submitting one or more gene identifiers. If an identifier is not unique, a window to resolve ambiguity will pop up before the network is fetched. To minimize ambiguities, the use of UniProt identifiers is recommended. After specifying the gene identifiers, users can select the desired search algorithm and adjust its parameters. To streamline the process, the app suggests default settings, which are the preferred configurations based on extensive testing and optimization. Available species are fetched from the API in order to enable up-to-date querying. In the *Subnetwork Selection* tab of the options panel, users can choose from a range of network expansion algorithms. The *Group gene search* algorithm is optimized for queries involving related genes. It expands the network by adding the N genes most strongly linked to the query genes. As opposed to this, the *Independent gene search* algorithm is well-suited for queries with unrelated genes, as it adds interactors independently to each query gene. The *MaxLink and TOPAS gene searches* were developed for identifying new disease gene candidates, and streamlining disease module identification. MaxLink evaluates direct connections between candidate and query genes, identifying statistically significant interactions to pinpoint potential therapeutic targets. TOPAS employs a top-down iterative approach that optimizes the identification of connected disease modules within complex networks.

Networks in the FunCoup website and the Cytoscape app share a common style definition. Nodes are coloured according to a species, and links are stylized depending on the type of the connection they represent, for example directed links are shown with arrow heads. Moreover, nodes and links each hold some additional information. Each node possesses a gene description, multiple aliases, and is associated with a list of tissues and pathways. Links on the other hand exhibit a positive predictive value (PPV), a Final Bayesian Score (FBS), the name of the supporting gold-standard network, as well as the summed Log likelihood ratio (LLR) scores for each of the ten evidence types and the 22 species from which evidence transfer via orthologs is done in FunCoup.

Additional link information can be fetched from the FunCoup API by picking a link and selecting the *Get Link Details* context menu item. As a result, a new window with information about the evidence of this link is opened. For each evidence, the corresponding LLR score and the respective supporting information is shown. Supplementary Figure 1 illustrates the link details between the proto-oncogene MYC and the Nucleophosmin NPM1. In this example, the interaction between MYC and NPM1 is predicted by FunCoup with high confidence, primarily identified as a directed regulatory interaction. This prediction is substantiated by robust evidence, including ChIP-Seq experiments targeting MYC in the MCF-7 human breast cancer cell line (ENCSR000DMM dataset, ENCFF377XCI bed file) and pertinent literature, such as a study demonstrating NPM1’s role in forming a complex with MYC to enhance the transcription of MYC target genes (Kim, Cho, and Park 2015).

Tissue specificity of gene expression can be a key factor in determining the context-dependent interactions between genes or proteins (Emig and Albrecht 2011). To enable such analysis, FunCoup 6 uses annotations from the Human Protein Atlas (v23.0) (Uhlén et al. 2015) and Bgee (v15.2) (Bastian et al. 2008) for a total of 1181 distinct tissues across 10 species with 131,258 unique genes with at least one tissue annotation. Tissue-filtered networks only display genes with non-zero expression and, for Bgee, a “gold quality” measure, using only tissues with at least 100 expressed genes. In addition to this, pathway annotations from KEGG (v108.1) contribute to a broader understanding, covering 453 distinct pathways across 22 species, representing 73,995 unique genes. The associated tissues and pathways of genes can be used to color and filter the network. This way a specific colour can be assigned to either each tissue or each pathway. Nodes without a colour or any association remain with the default color. Supplementary Figure 2A shows the resulting network of the query genes TSHR, PAX8 and SLC26A4, all linked to *thyroid hypoplasia*. The network shows which genes are active in the thyroid gland by coloring them. The network can be filtered in the *Filter* side panel by creating *Column Filters* (Supplementary Figure 2B). By providing a target column, the tissue/pathway name of interest and a condition (e.g., contains, equals, or RegEx), a subset of the nodes is shown. Multiple *Column Filters* can be combined.

The integration of tools like FunCoup and ClusterMaker2 (Utriainen and Morris 2023) in Cytoscape significantly enhances the analytical capabilities available to researchers, allowing for more customized and detailed network analyses. To illustrate this, Figure 1B shows an example of disease module prediction by TOPAS (Buzzao et al. 2022) using FunCoup with a link cutoff of PPV≥0.9 and a maximum number of connectors of 2. For this analysis, we queried FunCoup with all ORPHANET genes associated with *Myasthenia gravis*, which can be subcategorized into two distinct genetic origins: autoimmune and congenital. Despite their differences, both forms lead to neuromuscular dysfunction, and by integrating these gene sets, we explore potential shared mechanisms that may contribute to the understanding of the disease. The TOPAS module is composed of 37 genes, including 31 seeds and 6 connectors, with 63 links of which 39 are unknown, i.e. not in a gold standard in FunCoup. To facilitate the understanding of the disease mechanisms, we used Infomap (Rosvall and Bergstrom 2008) from ClusterMaker2 (v2.3.4) which retrieved six clusters. Many of these clusters correspond well to ORPHANET gene sets (Weinreich et al. 2008), indicating complex interactions among them (Supplementary Table 2). For instance, one cluster (purple) predominantly includes genes related to “Congenital myasthenic syndromes with glycosylation defect” and another one (green) to “Postsynaptic congenital myasthenic syndromes”.

Except for one case (yellow to green), the clusters are linked by connector genes added by TOPAS to make a single network module, suggesting that these connector genes are central to common mechanisms relevant to the disease. Among the connector genes, CREB1 is known to regulate the expression of genes involved in neural plasticity, survival and metabolism (Sakamoto, Karelina, and Obrietan 2011), and BHLHE40 is known to influence immune responses and inflammation (Cook et al. 2020). Given that *Myasthenia gravis* is a neuromuscular disorder (Mishra and Varma 2023), the dysregulation of these genes may impact muscle contraction and synaptic stability, potentially contributing to the symptoms of the disease which is characterized by muscle weakness due to impaired synaptic transmission at the neuromuscular junction. This demonstrates how advanced integrations can facilitate deeper insights into complex biological networks to reveal new relevant candidate disease genes and their interactions.

## DISCUSSION

FunCoup is a comprehensive resource for genome-wide functional association networks across 640 species, integrating multiple types of gene/protein interaction evidence using a unique bin-free redundancy-weighted Bayesian method. With the new FunCoup Cytoscape App, users familiar with the Cytoscape environment are provided with easy access to the FunCoup network resources. The Cytoscape environment further allows users to perform visualization and network analyses not available on the FunCoup website by using Cytoscape plugins and tools.

## DATA AVAILABILITY

The plugin is available as a JAR file at https://bitbucket.org/sonnhammergroup/funcoup_cytoscape/. The app is freely available for download from the Cytoscape app store https://apps.cytoscape.org/.

