## Supplementary Materials for "The FunCoup Cytoscape App: multi-species network analysis and visualization"

### MATERIALS AND METHODS

#### FunCoup

##### The database, the website and the API

FunCoup 6 is a comprehensive resource of genome-wide functional association networks. It employs a unique redundancy-weighted naïve Bayesian integration method to combine direct and indirect gene interactions with experimental data. FunCoup contrasts six gold standards against interactions expected by chance, to train Bayesian scoring functions for ten different types of evidence across seven broad categories: co-expression, co-localization, co-evolution, co-regulation, domain-domain interaction, genetic interaction, and protein-protein interaction. In FunCoup 6 networks for all 640 species in InParanoiDB 9 (Persson and Sonnhammer 2023) can be queried. Functional association networks for 23 species are extracted using species-specific gold standard data together with an extensive use of evidence transfer between them via InParanoid orthologs. Networks for the remaining 618 species are transferred from the closest FunCoup species, using orthologs from InParanoid, determined by NCBI taxonomy (Schoch et al. 2020) or orthophylogram distance.

The website and the API in FunCoup 6 were developed in Python (v3.10.8) using the Django framework (v4.1.1). The API is built to handle incoming queries at multiple endpoints, all of which return a response in the JSON format after interacting with the FunCoup database. Supplementary Table 1 lists all endpoints and their functionality. The JSON output is used by the Cytoscape app to create a graphical representation of the network.

#### Network search algorithms

We implemented gene search algorithms to query the database, serving specific biological questions and helping users to generate new hypotheses. These algorithms prioritize identifying the most strongly connected genes to a query and can operate on both individual genes and groups of genes.

##### Group and independent search

The default search is *group search*. By querying the database with query genes as a group, the N genes most strongly connected to the entire query set are added. Genes that interact with multiple query genes are more likely to be selected when the option to *prioritize common neighbors* is activated. As opposed to that, by querying the database for *each gene independently*, the N most strongly connected genes are added at each expansion step for each query gene.

##### MaxLink and TOPAS search

The Maxlink algorithm identifies candidate genes directly connected to query genes. It scores candidates based on their connections to the query genes and uses a hypergeometric test to find candidates whose interaction to the query set relative to other genes is significant (by default at p-value < 0.05).

The TOP-down Attachment of Seeds (TOPAS) algorithm iteratively connects the largest possible number of query genes in a top-down fashion. It preprocesses the network to ensure that seed genes are not in disconnected subnetworks. Potential connector genes lie on the shortest path (maximum distance of 3) between any two seed genes. Connector selection occurs during network pruning, where each gene's probability to be selected increases with the flow from a Random Walk with Restart (RWR) (Can, Çamoğlu, and Singh 2005) with a restart probability of 0.75.

##### Comparative interactomics search

The comparative interactomics feature is unique to FunCoup and allows aligning networks in multiple species. In release 6 this feature supports two search strategies to perform the query across the selected species. The previously implemented method is to perform the network search in the primary species only, and then find subnetworks in other species of only orthologs to genes in the found network. Orthology information is extracted from InParanoiDB 9. The new method is to perform a network search in each species network independently using orthologs to the query genes. This method will generally result in less network conservation than the previous method. As previously , by default only species-specific evidence is used for this analysis to avoid coherent network connections caused by orthology transfer.

#### Human vs SARS-CoV-2 search

In FunCoup, SARS-CoV-2 was added as an extension of the human interactome, incorporating the latest UniProt and IrefIndex data. This update, which is also accessible through the Cytoscape app, includes 2,879 links between viral and human proteins.

### Implementation of the App

The FunCoup App was developed in Java OpenJDK 17, using the OSGI framework (v6.0.0) and Cytoscape (v3.10.2). The development was supported by the build tool Maven (v3.8.1). UI components were implemented using the Java Swing GUI toolkit.

The FunCoup Cytoscape App queries the FunCoup 6 API. The main feature of fetching a network from the FunCoup API, results in a JSON list of nodes and links. These entries are parsed and converted to native structures in Cytoscape.

The default styling of the network is chosen to mirror the look of the FunCoup website. Nodes and links have customizable properties like fill color, shape, and size for nodes, and color, line type, and arrow shapes for links. The API outputs JSON files containing node colors, link directions, and other properties for visualization. Node colors are applied directly, link directions are visualized with arrow shapes, and node size is based on their degree. The maximum degree varies between networks based on the parameters used. Therefore, for each incoming network, a new continuous mapping is created. The actual node degree is translated to width and height values via a linear equation, with limits set by variables in the code.

### SUPPLEMENTARY FIGURES

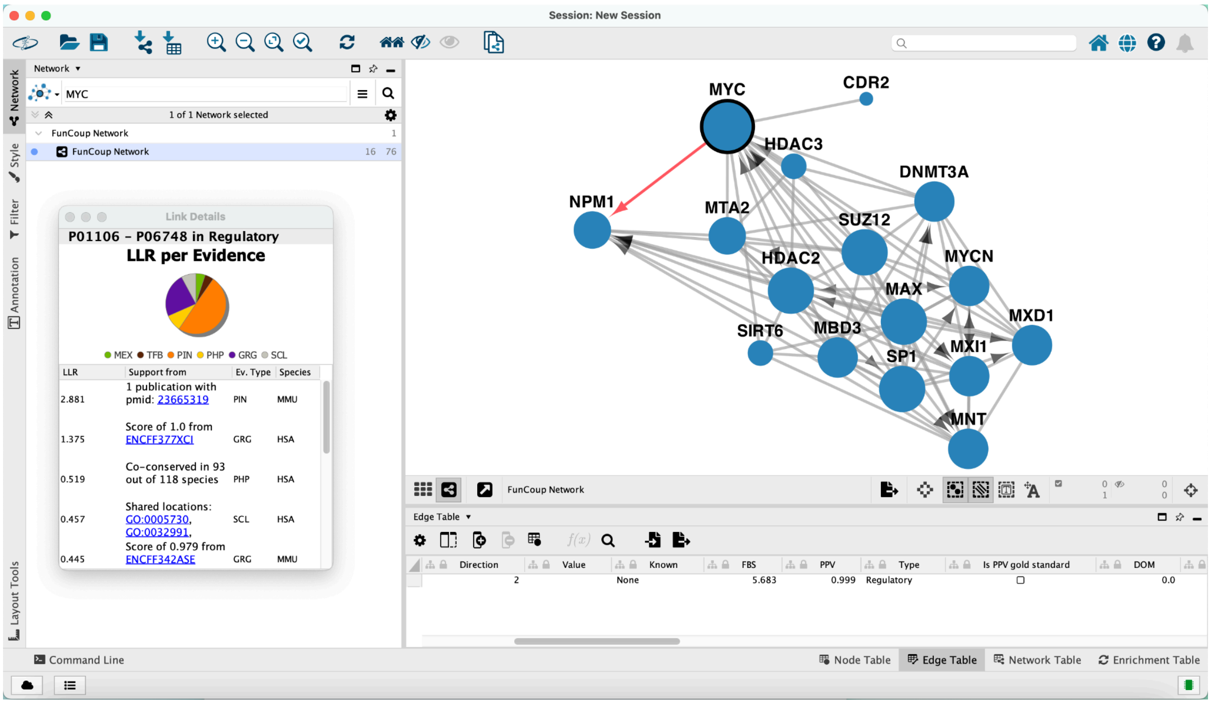


**Supplementary Figure 1. Details for the link between MYC & NPM1.** By selecting a link and clicking the corresponding context menu item, additional information can be fetched from the FunCoup API (link). As shown in the edge table, the link between MYC & NPM1 is of regulatory nature. MYC and NPM1 form a complex that enhances expression of MYC targets.

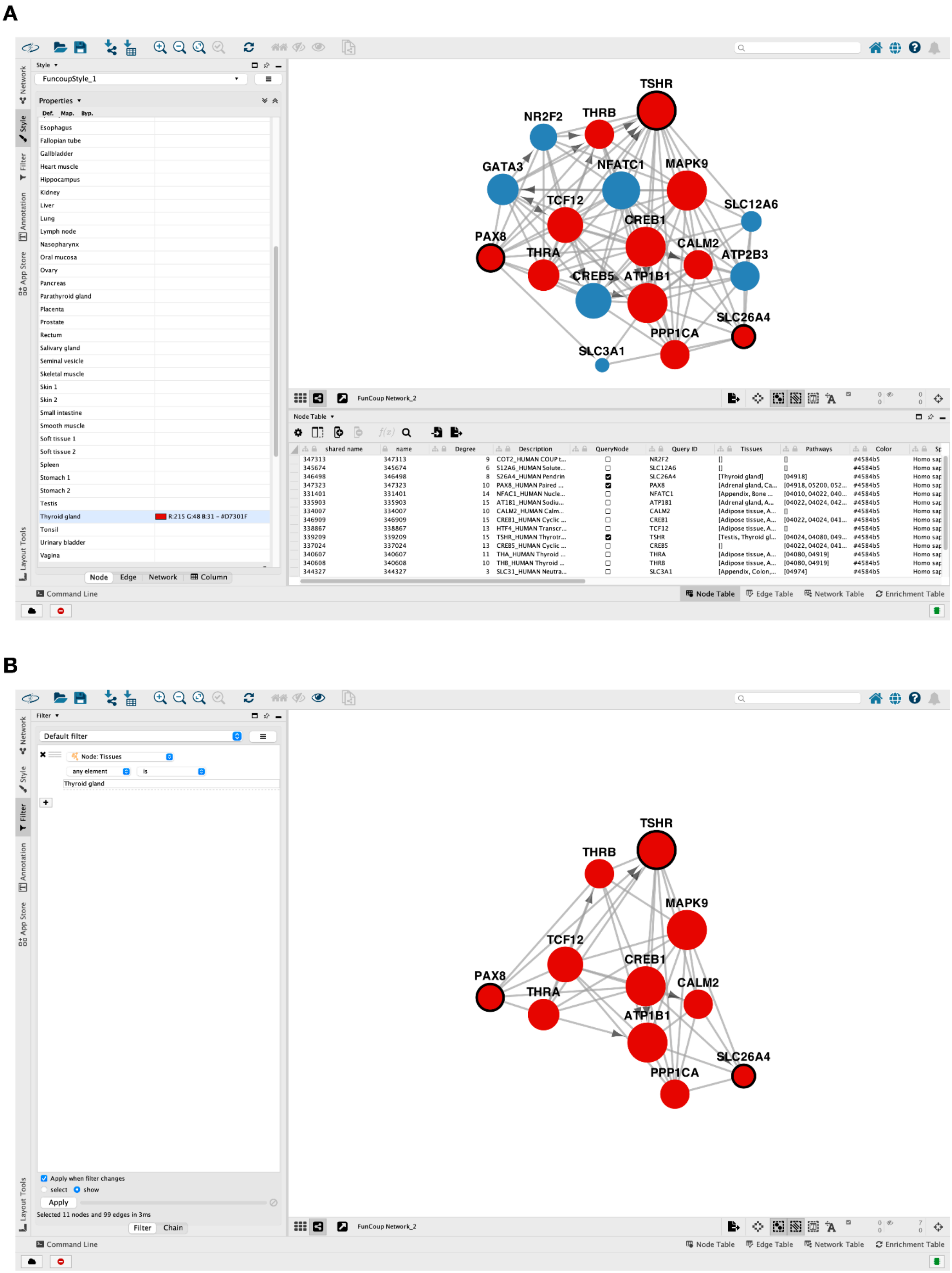


**Supplementary Figure 2. Example of coloring and filtering nodes by tissue.** The FunCoup API is queried for the genes TSHR, PAX8 and SLC26A4, all of which are associated with the disease *thyroid hypoplasia* (link). (A) Nodes with the *Thyroid gland* tissue annotation are coloured in red. (B) Nodes without the *Thyroid gland* tissue annotation are filtered out.

### SUPPLEMENTARY TABLES

**Supplementary Table 1 - List of FunCoup API Endpoints.** The FunCoup API offers four JSON endpoints. The Species endpoint is called at startup of the plugin and updates the available species in the Cytoscape app. Gene, Network and Link Details endpoints are called by user interaction and require additional information.

| API Endpoint | Functionality |
|---|---|
| **api/json/species** | Fetch a list of all the available species. No parameters have to be set for this query. |
| **api/json/gene** | Verify whether the query exists in the database and resolve any potential ambiguities. |
| **api/json/network** | Retrieve a network. For proper performance, the query parameters have to follow a certain specification. |
| **api/json/linkdetails** | Fetch supporting information for a link. Source, target and type of the link need to be specified in the query parameters. |

**Supplementary Table 2 - ORPHANET gene sets related to *Myasthenia gravis*.** The genes are 31 in total: AGRN, AK9, ALG14, ALG2, CHAT, CHRNA1, CHRNB1, CHRND, CHRNE, COL13A1, COLQ, CTLA4, DOK7, DPAGT1, GFPT1, GMPPB, HLA-DQA1, LAMB2, LRP4, MUSK, MYO9A, RAPSN, RYR1, SCN4A, SLC18A3, SLC25A1, SLC5A7, SNAP25, SYT2, TNFRSF11A, VAMP1. AGRN and COL13A1 are present both in *Postsynaptic* and *Presynaptic congenital myasthenic syndromes*.

| ORPHANET gene set | Gene Symbols |
|---|---|
| Adult-onset myasthenia gravis | TNFRSF11A, CTLA4, HLA-DQA1 |
| Congenital myasthenic syndromes with glycosylation defect | ALG2, DPAGT1, GFPT1, ALG14, GMPPB |
| Congenital myopathy with myasthenic-like onset | RYR1 |
| Postsynaptic congenital myasthenic syndromes | RAPSN, SCN4A, CHRNA1, CHRNB1, CHRND, CHRNE, DOK7, MUSK, LRP4, AGRN, AK9, COL13A1 |
| Presynaptic congenital myasthenic syndromes | CHAT, AGRN, SLC25A1, SNAP25, SYT2, MYO9A, SLC5A7, VAMP1, SLC18A3, COL13A1 |
| Synaptic congenital myasthenic syndromes | COLQ, LAMB2 |
